# IPMK modulates insulin-mediated suppression of hepatic glucose production

**DOI:** 10.1101/2022.03.10.483655

**Authors:** Ik-Rak Jung, Frederick Anokye-Danso, Sunghee Jin, Rexford S. Ahima, Sangwon F. Kim

**Author notes:** **Corresponding Authors:** Sangwon F. Kim, PhD and Rexford S. Ahima, MD, PhD, Department of Medicine, Division of Endocrinology, Diabetes, and Metabolism, Johns Hopkins University, 5501 Hopkins Bayview Cir, AAC 2A44/62, Baltimore, Maryland, USA. 21224.

## Abstract

Hepatic glucose production is crucial for the maintenance of normal glucose homeostasis. Although hepatic insulin resistance contributes to excessive glucose production, its mechanism is not well understood. Here, we show that inositol polyphosphate multikinase (IPMK), a key enzyme in inositol polyphosphate biosynthesis, plays a role in regulating hepatic insulin signaling and gluconeogenesis both in vitro and in vivo.IPMK-deficient hepatocytes exhibit decreased insulin-induced activation of Akt-FoxO1 signaling. The expression of mRNA levels of phosphoenolpyruvate carboxykinase 1 (Pck1) and glucose 6-phosphatase (G6pc), key enzymes mediating gluconeogenesis, are increased in IPMK-deficient hepatocytes compared to wild type (WT) hepatocytes. Importantly, re-expressing IPMK restores insulin sensitivity and alleviates glucose production in IPMK-deficient hepatocytes. Moreover, hepatocyte-specific IPMK deletion exacerbates hyperglycemia and insulin sensitivity in mice fed a high-fat diet (HFD), accompanied by an increase in hepatic glucose production during pyruvate tolerance test and reduction in Akt phosphorylation in IPMK deficient liver. Our results demonstrate that IPMK mediates insulin signaling and gluconeogenesis and may be potentially targeted for treatment of diabetes.

**Highlights:** IPMK expression is reduced in livers of HFD-fed mice.

Hepatocyte-specific deletion of IPMK in mice aggravated HFD-induced insulin resistance.

Loss of IPMK decreased insulin-induced activation of Akt-FoxO1 signaling, leading to the increase of glucose production in hepatocytes.

## 1. Introduction

IPMK is an enzyme essential for converting inositol polyphosphate (IP)3 to IP4, and IP4 to IP5 in mammalian cells [1]. In addition to its primary role in generating IP5, IPMK also possesses phosphatidylinositol 3 kinase (PI3K) activity that phosphorylates phosphatidylinositol 4,5-bisphosphates (PIP2) to produce phosphatidylinositol 3,4,5-trisphosphate (PIP3) which is proposed to contribute to activation of Akt/PKB upon exposure to various hormones or growth factors such as insulin [2; 3]. IPMK is also regulates cellular signaling in a catalytic independent manner. For example, IPMK physically binds to AMP-activated protein kinase (AMPK) which is a central regulator of energy homeostasis, modulating cellular glucose homeostasis by enhancing its downstream signaling [4; 5]. Moreover, IPMK interacts with mammalian target of rapamycin (mTOR) to stabilize the mTOR-raptor complex and enhances protein synthesis [6]. Furthermore, IPMK plays a role in gene expression as a transcriptional coactivator. IPMK binds serum response factor (SRF) and stabilizes SRF binding to its promoter, serum response element (SRE) or directly interact with CREB-binding protein (CBP) to induce a wide range of immediate early gens (IEGs), such as c-jun and c-fos [7; 8]. However, very few studies have elucidated the physiological roles of IPMK on energy metabolism at the systemic level in animal models. Recently, it has been reported that adipocyte IPMK is dispensable for normal adipose functions and whole-body metabolism [9].

The liver plays a central role in the regulation of glucose homeostasis. In particular, the liver takes up glucose and synthesizes glycogen to prevent postprandial hyperglycemia. On the other hand, in the fasted state, the liver supplies glucose through hepatic glucose production from gluconeogenesis and glycogenolysis to maintain normal glucose levels. These processes are highly regulated by hormones such as glucagon and insulin produced by pancreatic islets. Glucagon is a primary regulator of hepatic glucose production (HGP) during fasting while insulin suppresses glucose production and clears plasma glucose in the postprandial state [10; 11]. Insulin directly acts on hepatocytes to suppress HGP via multiple mechanisms, including changes in substrate metabolism and gene expression [12; 13]. Mechanistically, insulin regulates hepatic gluconeogenic gene expression by inhibiting transcription related to gluconeogenesis such as G6pc and Pck1. It has been suggested that insulin regulation of HGP in the liver is primarily modulated via pAkt-mediated inactivation of glycogen synthase kinase 3ß (GSK3) and FoxO1 to regulate glycogenolysis and gluconeogenic gene expression, respectively [14]. However, in type 2 diabetic conditions, insulin fails to regulate hepatic glucose production, leading to hyperglycemia [15; 16].

Various studies have examined the molecular mechanisms by which insulin fails to suppress HGP [17]. Here, we investigate a role of IPMK in the control of hepatic glucose production in mice with a hepatocyte-specific IPMK deficiency and identify IPMK as a regulator of gluconeogenesis. We show that loss of hepatic IPMK decreases insulin-stimulated Akt phosphorylation and its downstream signaling pathway, ultimately enhancing glucose production during the refeeding state. Moreover, a loss of hepatic IPMK exacerbates HFD-induced glucose intolerance and insulin resistance. We elucidate a role of hepatic IPMK in glucose metabolism through regulation of Akt phosphorylation and gluconeogenesis-related gene expression.

## 2. Materials and Methods

### 2.1. Animals

Experiments were performed in accordance with the Institutional Animal Care and Use Committee guidelines with its approvals. The LKO mice were generated by crossing Albumin-CreTg/+ mice (Jackson Laboratories) with mice homozygous for a “floxed” exon 6 of IPMK (IPMK *fl//fl*). The control mice for this study were IPMK *fl/fl* (WT) mice. Mice were housed under standard conditions in a temperature- and humidity-controlled facility with a light–dark cycle of 12 h (lights on at 07:00) and fed ad libitum at the Johns Hopkins University (Baltimore, MD). WT and LKO mice aged 6 wk were fed a ND (containing 13% fat, 23.3% protein, and 63.7% carbohydrate; LabDiet, St. Louis, MO, USA) or HFD (containing 60 kcal% fat, 20% protein, and 20% carbohydrate; Research Diets, New Brunswick, NJ, USA) for 12 wk. Blood was collected from the tail after 5 h fasting (08:00 AM – 1:00 PM). Blood glucose levels were measured using Contour glucometer (Ascensia Diabetes Care, Parsippany, NJ, USA). Plasma insulin was measured using the mouse Insulin ELISA Kit (Crystal Chem, Elk Grove Village, IL, USA) according to the manufacturer’s protocol.

### 2.2. Mouse metabolic studies

The body composition was measured using ^1^H Magnetic Resonance Spectroscopy (Echo MRI, Houston, TX). To assess energy homeostasis, the mice were housed singly in in a Comprehensive Laboratory Animal Monitoring System (CLAMS, Columbus Instruments, Columbus, OH) on a 12-hour light/dark cycle (lights on at 07:00) and cage temperature was maintained at 22°C. After 3 days of habituation, food intake, energy expenditure and locomotor activity were continuously monitored for 2 days [18].

Glucose homeostasis as assessed using glucose tolerance test (GTT), insulin tolerance test (ITT) and pyruvate tolerance test (PTT). For GTT, the mice were fasted for 5 hours in the morning and glucose solution (1 g/kg body weight) was injected intraperitoneally. Tail blood glucose was measured at time 0, 15, 30, 60 and 120 minutes using a Contour glucometer (Ascensia Diabetes Care, Parsippany, NJ, USA). For ITT, mice were fasted or 5 hours in the morning, regular human insulin (Humulin 1 U/kg body weight) was injected IP, and tail blood glucose was measured at time 0, 15, 30, 60 and 120 minutes. For PTT, the mice were fasted overnight, sodium pyruvate (2 g/kg body weight) was injected IP, and tail blood glucose was measured at time 0, 15, 30, 60 and 120 minutes [19].

### 2.3. In vivo insulin signaling

Mice on HFD were fasted for 5 hours, anesthetized and the portal vein was exposed. Human insulin (2U, Humulin R; Eli Lilly, Indianapolis, IN, USA) (100 μl) was injected into the portal vein. Liver tissues were harvested 3 min, and snap frozen for immunoblot analysis [20].

### 2.4. Liver glycogen content measurement

Fed or overnight fasted mice were euthanized, liver samples rapidly harvested and snap frozen and glycogen levels were measured using Glycogen Colorimetric Assay Kit (Biovision, Milpitas, CA, USA).

### 2.5. Cell Culture

Primary hepatocytes were isolated from WT and LKO male mice, using collagenase perfusion followed by filtration through a 100 μm mesh as previously described [21]. Cells were all maintained in Media 199 (Corning Inc, Corning, NY, USA) supplemented with 10% fetal bovine serum (MilliporeSigma, Burlington, MA, USA) and 100 units/mL penicillin/streptomycin (MilliporeSigma, Burlington, MA, USA). Transfection of primary hepatocytes was performed by using lipofectamine 3000 transfection reagent (Thermo Fisher Scientific, Waltham, MA, USA), according to the protocol of the manufacturer.

### 2.6. Hepatocyte glucose production assay

Primary mouse hepatocytes were treated with 100 nM insulin (MilliporeSigma, Burlington, MA, USA) for 3 h in glucose-free DMEM (Thermo Fisher Scientific, Waltham, MA, USA) supplemented with 20 mM sodium lactate (MilliporeSigma, Burlington, MA, USA), 2 mM sodium pyruvate (MilliporeSigma, Burlington, MA, USA), 2 mM L-glutamine (Corning Inc, Corning, NY, USA) and 15 mM HEPES (Thermo Fisher Scientific, Waltham, MA, USA). The glucose production in the culture medium was determined using the Autokit Glucose (FIJIFILM Wako Pure Chemical Corporation, Osaka, Japan).

### 2.7. Immunoblotting

Samples were lysed in RIPA buffer containing inhibitors and heated at 95°C for 5 min prior to electrophoresis. Proteins were transferred to a 0.2-mm nitrocellulose membrane, blocked with 5% nonfat dry milk, and incubated with primary antibodies at 4°C overnight. Immunoblotting was conducted with the following antibodies: IPMK from Novus Biologicals and pAkt-S473, pAkt-T308, Akt, pFoxO1, FoxO1, pGSK3ß-S9, GSK3ß, GAPDH from Cell Signaling Technology. Blots were imaged and quantitated using an Odyssey Near-Infrared Scanner (Li-Cor Biosciences, Lincoln, NE, USA).

### 2.8. Quantitative real-time PCR

Total RNA was isolated from WT or LKO mouse liver tissue or primary hepatocytes using TRIZol reagent (Thermo Fisher Scientific, Waltham, MA, USA), and chloroform (MilliporeSigma, Burlington, MA, USA), precipitated in 2-propanol (MilliporeSigma, Burlington, MA, USA), washed in ethanol (MilliporeSigma, Burlington, MA, USA), and quantified using Epoch Microplate Spectrophotometer (Agilent Technologies, Santa Clara, CA, USA). mRNA was reverse transcribed into cDNA using ProtoScript^®^ II First Strand cDNA Synthesis Kit (BioLabs, Ipswich, MA, USA). Ribosomal Protein L32 (RPL32) mRNA was used as the invariant endogenous control and melting curve analysis was run to verify specificity of each amplicon. All reactions were performed in triplicate. The relative amounts of the RNAs were calculated using the comparative threshold cycle method. Primers were as follows: Pck1 forward 5’-CATAACGGTCTGGACTTCTCTGC-3’, reverse 5’-GAATGGGATGACATACATGGTGCG-3’; G6pc forward 5’-ACACCGACTACTACAGCAACAG-3’, reverse 5’-CCTCGAAAGATAGCAAGAGTAG-3’; Rpl32 forward 5’-AAGCGAAACTGGCGGAAACC-3’, reverse 5’-CCCATAACCGATGTTGGGCA-3’.

### 2.9. Statistical Analysis

All statistical analyses were performed using GraphPad Prism 5.0 (GraphPad, San Diego, CA, USA), and the data are presented as mean ± SEM. Comparisons between two groups were done using Student’s t test. Comparisons among multiple groups were done using ANOVA. The threshold for statistical significance was set at P < 0.05, and Bonferroni’s multiple-comparison test was used for post hoc analyses.

## 3. Results

### 3.1. Liver-Specific Knockout of IPMK Disrupts Insulin Signaling in Mice

IPMK is highly expressed in the liver of mice fed a normal diet (ND) and decreased in mice fed a high fat diet (HFD) (Supplementary Fig. 1A and B). To investigate the function of IPMK, we generated liver specific IPMK deleted mice (LKO) by crossing *Ipmk^f/f^* (neomycin deleted) [2] mice with Albumin-Cre mice. IPMK protein was completely deleted in LKO liver tissue but not in other tissues such as muscle and adipose tissues (Supplementary Fig. 2B-D). LKO mice displayed no difference in body weight (Supplementary Fig. 2E) and the liver to body weight ratio was not affected (Supplementary Fig. 2F). To understand the in vivo role of IPMK in hepatic insulin signaling, we characterized glucose metabolism and insulin signaling in LKO versus control (WT) mice. Blood glucose concentration was similar between LKO and WT in the fasted state but significantly higher in LKO than WT after refeeding (Fig. 1A). Plasma insulin concentrations did not differ between LKO and WT mice in both fasted and refed conditions (Fig. 1B). There were no significant differences in glycogen content were observed between LKO and WT mice in the fasted state (Fig. 1C) whereas LKO mice had reduced liver glycogen in the refed state compared with WT mice (Fig. 1C). Since blood glucose and liver glycogen contents were altered without any changes in plasma insulin concentrations in the refed state, we investigated the hepatic insulin response which is the key signaling pathway that regulates hepatic glucose production. Following refeeding, the pAkt level tended to decrease in LKO mice than in WT (Fig 1D). Moreover, the phosphorylated levels of GSK3ß (pGSK3ß) and FoxO1 (pFoxO1) were reduced in LKO mice (Fig. 1D). The mRNA expression of Pck1 and G6pc were not suppressed in LKO as much as in WT mice in the refed state, showing 8- and 2.6-fold higher than WT (Fig. 1E). Together, these data suggest that a loss of IPMK attenuates hepatic insulin signaling leading to higher glucose levels in refed LKO mice.

**Figure 1.**
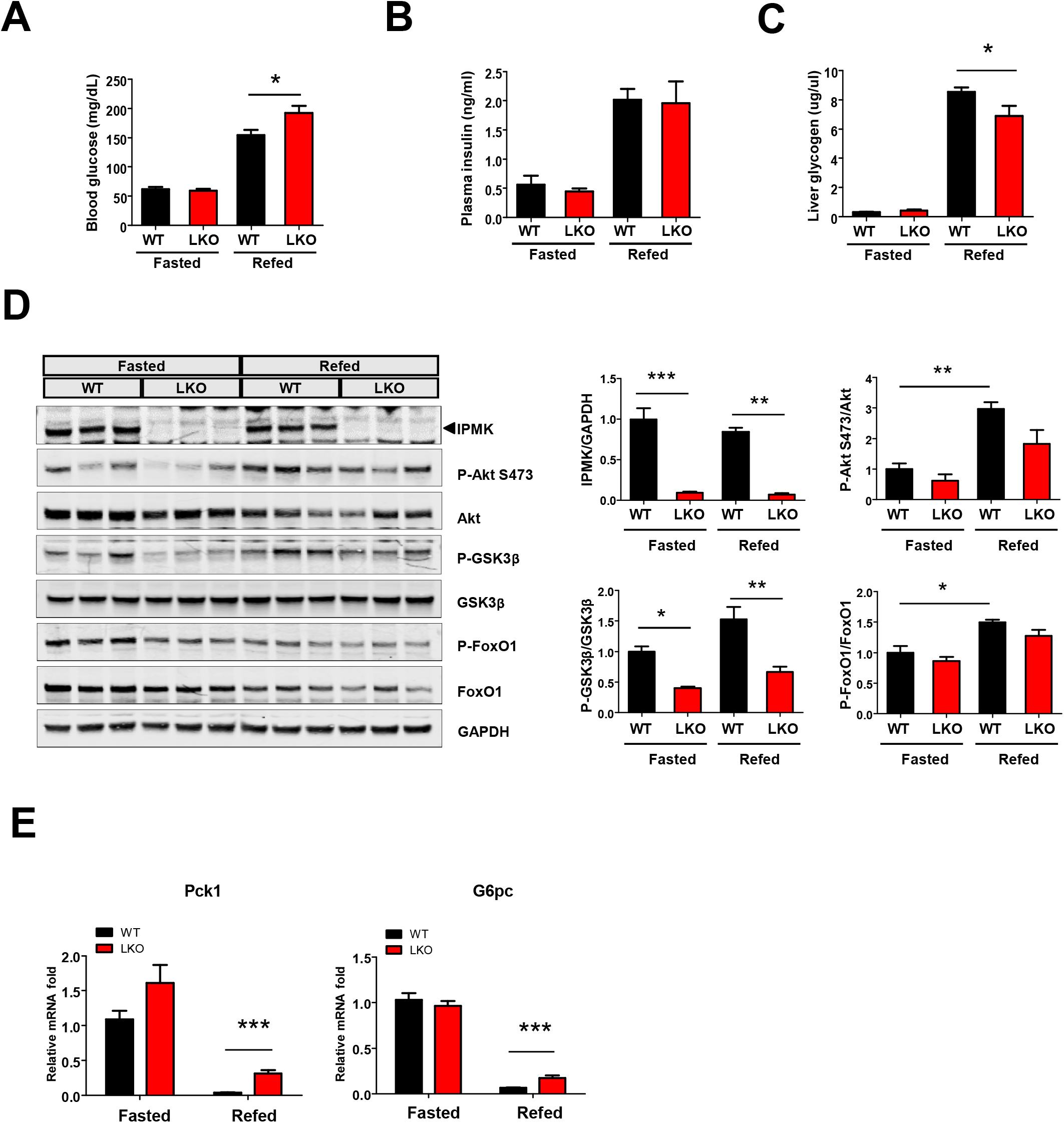
Effects of hepatic IPMK deficiency in fasted versus refed states. (A) Blood glucose, (B) plasma insulin and (C) liver glycogen levels were measured in male WT and LKO mice after overnight fasting (WT, n = 5; LKO, n = 5-8) or overnight fasting and 4 hours refeeding (WT, n = 5; LKO, n = 5-8). (D) Immunoblotting for insulin signaling molecules and (E) mRNA levels of gluconeogenic genes in the liver from WT and LKO mice after overnight fasting or overnight fasting and 4 hours refeeding (D; n = 3, E; n = 4 per group). Levels of indicated proteins were quantified. Data are mean ± SEM; *p < 0.05, **p < 0.01, ***p < 0.001

### 3.2. Loss of hepatic IPMK alters body composition and energy homeostasis in HFD-fed mice

To better understand the role of hepatic IPMK in diet-induced obesity, both male and female mice (WT and LKO) were fed a normal diet (ND) versus a high fat diet (HFD) for 12 weeks. On ND, there was no difference in body weights of WT and LKO male mice (Fig. 2A). However, the fat mass was slightly increased in LKO male mice compared to WT, while the lean mass did not differ under ND (Fig. 2B). In contrast, the body weights of HFD LKO male mice were significantly higher than control mice after 7 weeks on HFD (Fig. 2C). Moreover, HFD feeding led to a significant increase in fat mass in LKO male mice, accompanied by a decrease in lean mass compared to WT mice (Fig. 2D). In contrast, hepatic IPMK deletion did not affect body weight or body composition in female mice fed either ND or HFD (Supplementary Fig. 3A and B). We assessed the effects of hepatic IPMK deletion on energy homeostasis (Fig. 2E-J). WT and LKO male mice fed ND displayed similar levels of food intake and energy expenditure (Fig. 2E and G). On the HFD, the LKO mice consumed a similar amount of food as WT mice (Fig. 2H), but the slope of the energy expenditure versus lean mass graph trended lower in LKO compared to WT mice (P=0.09) (Fig. 2J). The respiratory exchange ratio (RER) was similar in both genotypes (Fig. 2I).

**Figure 2.**
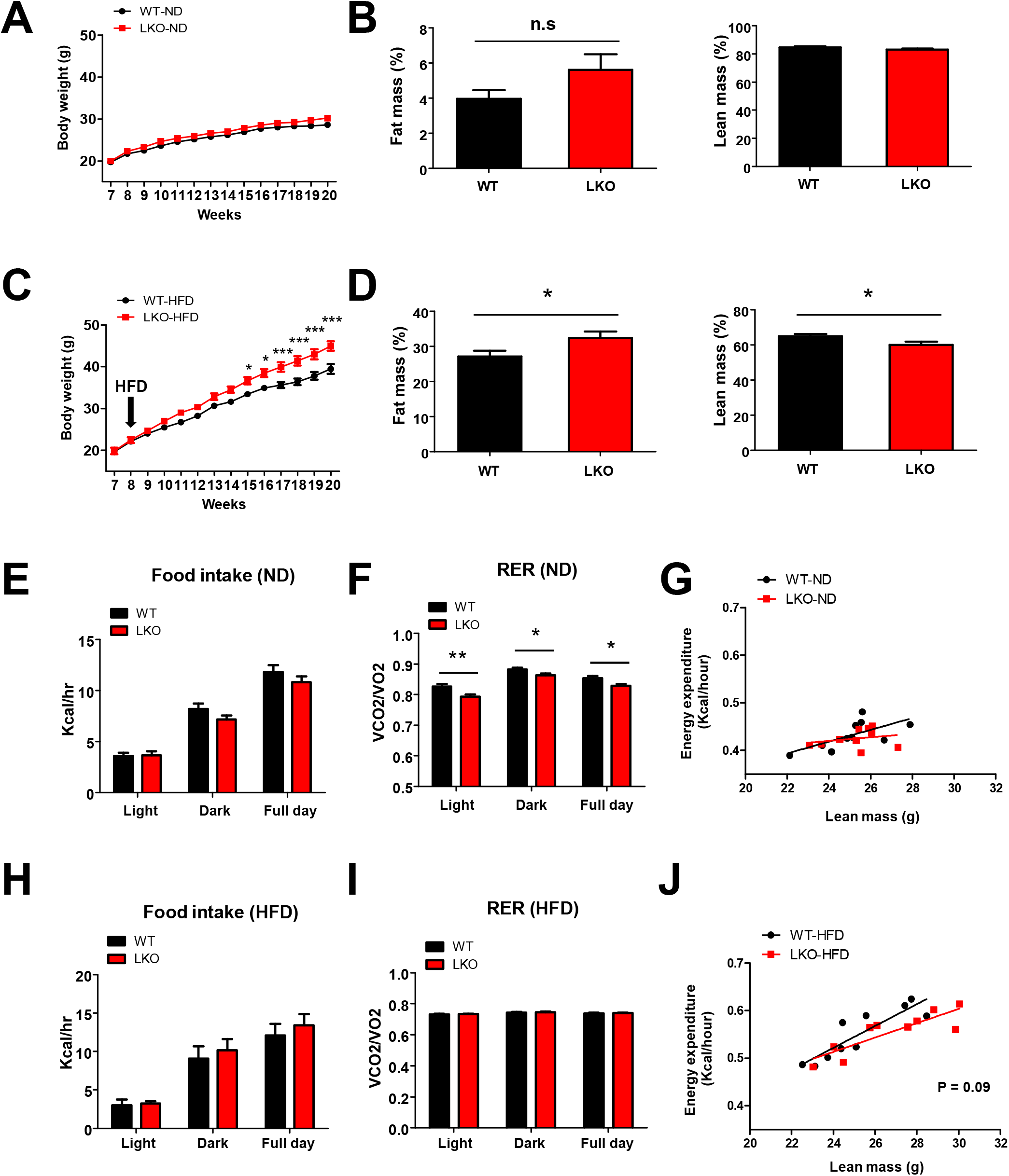
Effects of hepatic IPMK deficiency on energy homeostasis. Body weight of WT and LKO male mice fed (A) ND (WT; LKO, n = 10) or (C) HFD (WT= 10; LKO, n = 8) was measured weekly from age 8 to 20 weeks. Lean and fat mass was determined by ^1^H-MRS analysis of 20-week-old mice fed (B) ND (WT; LKO, n =10) or (D) HFD (WT, n = 10; LKO, n = 8). Indirect calorimetry was performed after 12 weeks of ND or HFD. Mean light-, dark and full day food intake (kcal/h) and RER (VCO_2_/VO_2_) of WT and LKO mice fed (E and F) ND (WT; LKO, n = 10) or (H and I) HFD (WT, n = 10; LKO, n = 8) were measured. Regression analysis of energy expenditure against lean mass was measured in mice fed (G) ND (WT, n = 10, LKO n =10) or (J) HFD (WT, n = 10; LKO, n = 8). The data are mean ± SEM; *p < 0.05, **p < 0.01, ***p < 0.001.

### 3.3. Loss of hepatic IPMK increases glucose and blunts insulin sensitivity

Glucose homeostasis was evaluated using intraperitoneal glucose tolerance test (GTT), insulin tolerance test (ITT) and pyruvate tolerance test (PTT). LKO and WT mice fed ND showed similar patterns of GTT and ITT (Fig. 3A and B). On HFD, LKO mice exhibited significantly higher glucose levels during GTT and ITT compared to WT mice (Fig. 3A and B). Furthermore, we determined the hepatic glucose production in vivo by performing PTT. Administration of pyruvate, a gluconeogenic substrate, similarly increased the blood glucose levels in ND WT and LKO mice (Fig. 3C) but in the HFD-fed group, pyruvate-induced glucose production was significantly higher in LKO mice compared to WT mice (Fig. 3C). A loss of hepatic IPMK in female mice did not affect PTT (Supplementary Fig. 3C). Since LKO male mice showed impaired glucose tolerance and elevated glucose production on HFD, we examined insulin-stimulated pAkt in HFD-fed WT versus LKO male mice. Loss of hepatic IPMK decreased the basal levels of pAkt and the insulin-stimulated pAkt level was blunted (Fig 3D), suggesting that loss of IPMK disrupts insulin-mediated glucose regulation in the liver.

**Figure 3.**
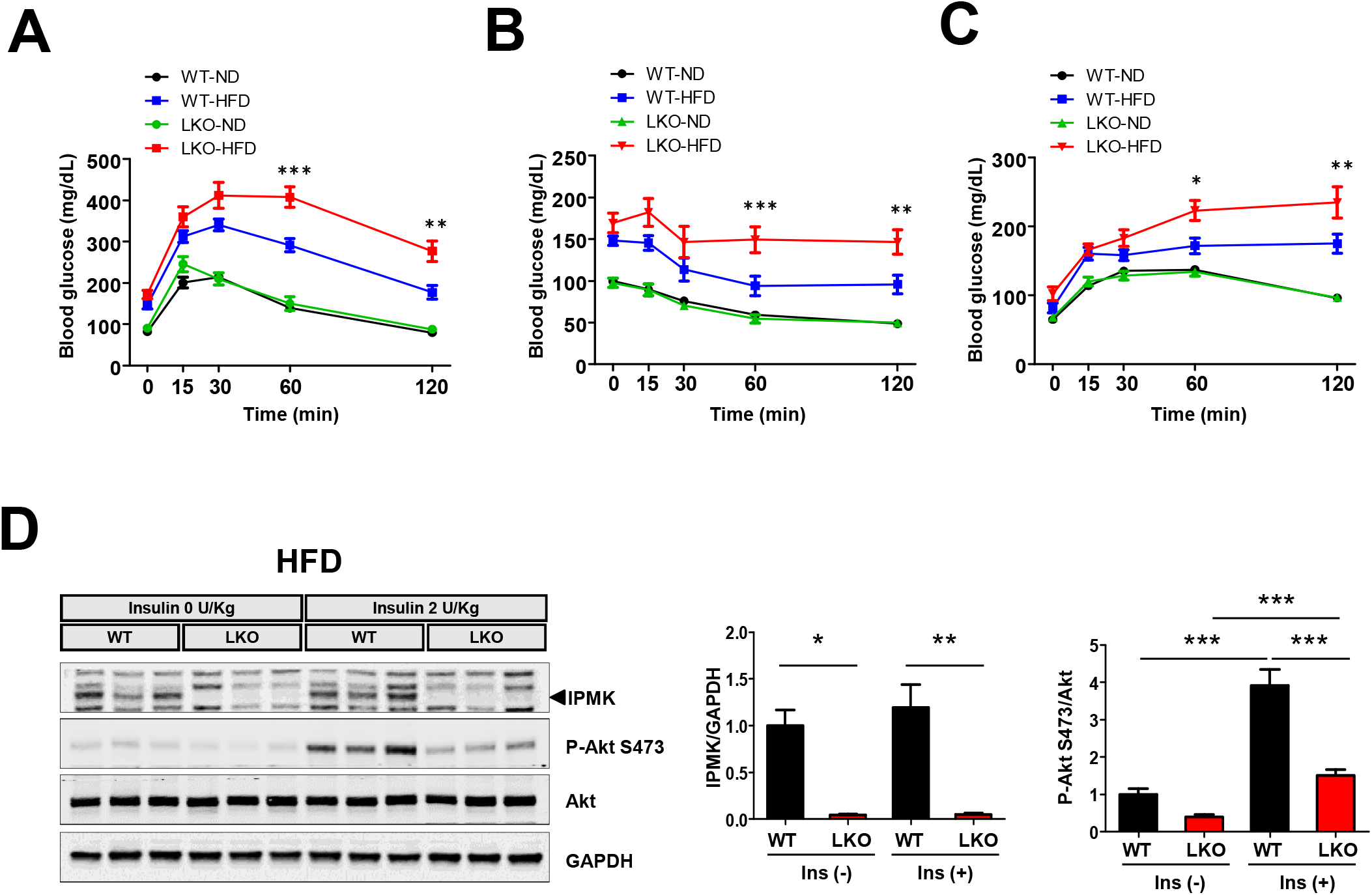
Effects of hepatic IPMK deficiency on glucose homeostasis. (A) GTT, (B) ITT and (C) PTT were performed in WT and LKO male mice (n = 8-10 per group). (D) Immunoblotting of pAkt after administering insulin via portal vein. Representative bands from each group (n = 5) are shown in immunoblotting images. Levels of indicated proteins were quantified. Data are mean ± SEM; *p < 0.05, **p < 0.01, ***p < 0.001.

### 3.4. Loss of IPMK Impairs Insulin-mediated Suppression of Hepatic Gluconeogenesis

To define the role of IPMK in hepatic metabolism, we examined insulin’s actions on hepatic gluconeogenesis and related gene expression in primary mouse hepatocytes. First, we investigated the insulin-stimulated Akt phosphorylation, a hallmark of insulin signaling. Insulin-mediated activation was significantly attenuated in hepatocytes from LKO mice compared to WT hepatocytes (Fig. 4A). One of the main actions of insulin in hepatocytes is the suppression of gluconeogenesis through activation of the Akt/Fork Head Box O1 (FoxO1) signaling pathway [12]. Consistent with its regulation of Akt signaling, Foxo1 phosphorylation was also reduced in LKO hepatocytes (Fig. 4A). Importantly, mRNA levels of Pck1 and G6pc, key gluconeogenic genes were increased in LKO hepatocytes compared with in WT upon insulin treatment (Fig. 4B and C). In addition, we examined glucose production after insulin treatment to determine whether the inhibitory effect of insulin on gluconeogenesis was indeed blunted in LKO hepatocytes. Insulin treatment reduced glucose production in WT hepatocytes, whereas the insulin-induced suppression was abolished in LKO hepatocytes (Fig. 4D). To further confirm the role of IPMK in hepatic gluconeogenesis, we rescued LKO hepatocytes by overexpressing IPMK. Overexpression of IPMK suppressed glucose production to normal levels (Fig. 4E). As predicted, insulin-stimulated Akt phosphorylation was increased to normal levels in LKO hepatocytes (Fig. 4F). Since the loss of IPMK decreased Akt phosphorylation and increased gluconeogenesis in primary hepatocytes, we examined whether Akt activation could suppress glucose production in IPMK-deficient hepatocytes. SC79, which is a specific Akt activator, enhances Akt phosphorylation and activation in various cell types [22]. Treatment with SC79 significantly reduced glucose production in LKO hepatocytes (Fig. 5A), accompanied by an increase in Akt phosphorylation (Fig. 5B) and reduction in the expression of gluconeogenic genes (Fig. 5C and D). These data indicate that hepatic IPMK plays an important role in insulin-mediated inhibition of gluconeogenesis.

**Figure 4.**
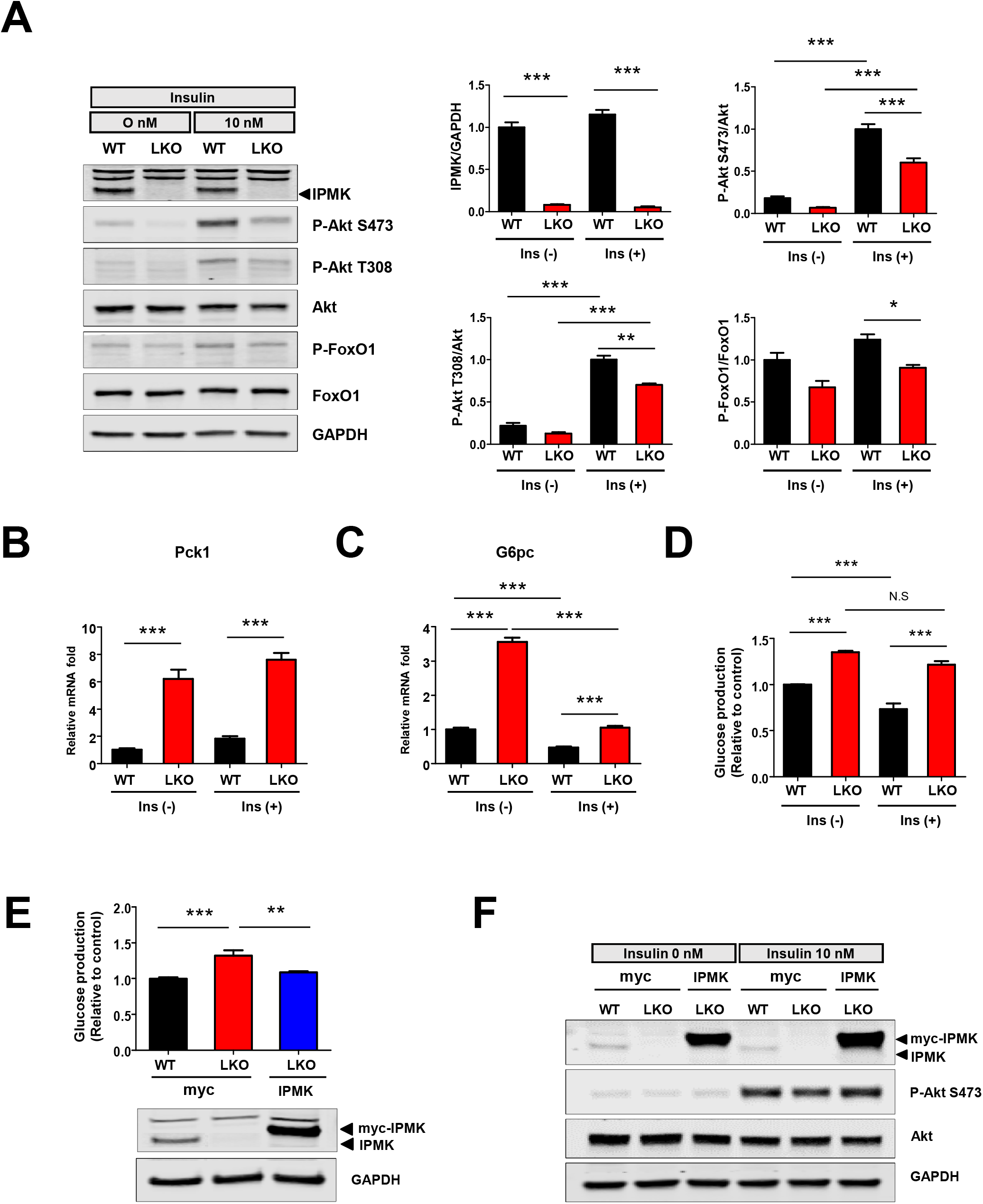
Loss of hepatic IPMK decreases hepatocellular insulin signaling and increases glucose production. (A) Primary mouse hepatocytes from WT and LKO mice were treated with 10 nM insulin for 10 minutes and cell lysates were subjected to immunoblotting. Levels of indicated proteins (phosphorylated vs total insulin signaling proteins, IPMK, GAPDH) were quantified. mRNA expression of gluconeogenic genes, (B) Pck1 and (C) G6pc was measured in WT and LKO primary hepatocytes after treatment with insulin (10 nM) for 4 hours by real-time quantitative PCR (qPCR). (D) Glucose production from lactate and pyruvate was determined by measuring the amount of glucose released from WT and LKO primary hepatocytes with and without insulin (10 nM) for 5 hours. (E) Primary hepatocytes from LKO mice were overexpressed with IPMK, and the glucose production was determined. Representative immunoblot images from at least 3 independent assays are presented. (F) Primary hepatocytes from LKO mouse were overexpressed with IPMK and treated with 10 nM insulin for 10 minutes and cell lysates were subjected to immunoblotting. The representative Western bolt images from at least 3 independent assays are presented. Data are mean ± SEM; *p < 0.05, **p < 0.01, ***p < 0.001, n = 3.

**Figure 5.**
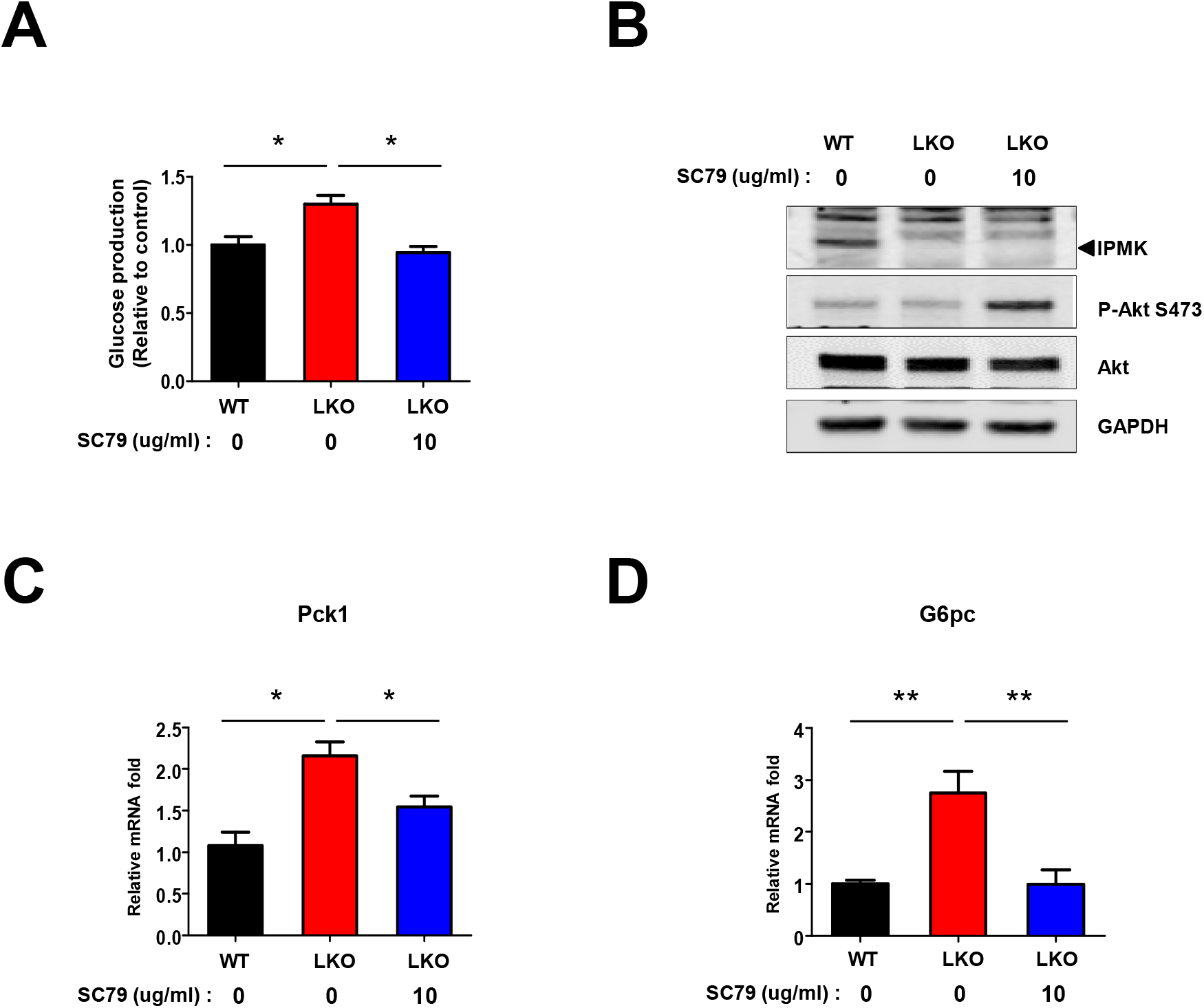
Activation of Akt decreases glucose production and gluconeogenic gene expression in LKO primary hepatocytes. (A) Glucose production from lactate and pyruvate was determined by measuring the amount of glucose released from WT and LKO primary hepatocytes with and without Akt activator treatment (SC79) for 5 hours. (B) Primary mouse hepatocytes from WT and LKO mice were treated with SC79 for 5 hours and cell lysates were subjected to immunoblotting. The representative immunoblot images from at least 3 independent assays are presented. mRNA expression of gluconeogenic genes, (C) Pck1 and (D) G6pc were measured in WT and LKO primary hepatocytes after treatment with SC79 for 5 hours by real-time quantitative PCR (qPCR). Data mean ± SEM; *p < 0.05, **p < 0.01, n = 3.

## 4. Discussion

There have been numerous reports describing the cellular roles of IPMK on metabolic pathways, but few studies have investigated the causal functions of IPMK utilizing tissue specific cells or animal models. In this study, we found that a loss of IPMK increased glucose production in primary mouse hepatocytes by increasing the expression of gluconeogenic genes via Akt-FoxO1 pathway. Hepatic IPMK deficiency increased body weight and fat in HFD male LKO mice, and this was accompanied by impaired glucose tolerance, insulin resistance and increased glucose production. IPMK is the rate-limiting enzyme in the synthesis of soluble inositol polyphosphates (IPs) and inositol pyrophosphates (PP-InsPs), and an important component in the recycling of phosphatidylinositol lipids [3]. A loss of IPMK resulted in 50% reduction in PIP3 in mouse embryonic fibroblasts (MEFs) [2]. Furthermore, PIP3-dependent Akt phosphorylation and activation of its downstream targets such as phospho-PRAS40, phospho-TSC2, phospho-FoxO1/3A and phospho-GSK3b are blunted in IPMK-deficient MEFs [2]. The Akt-FoxO1 pathway is critical for hepatic insulin-regulated glucose homeostasis [23]. Phosphorylation of FoxO1 by activated Akt was paralleled inhibition of Pck1 and G6pc mRNA expression [24] and FoxO1 loss of function in the liver reduces hepatic glucose production [25]. We have confirmed that hepatic IPMK can also regulate Akt phosphorylation as reported in other cell types [2; 26; 27]. In addition, a pharmacological activator of Akt, which is independent of PIP3 levels, suppressed gluconeogenesis in IPMK-deficient hepatocytes indicating that IPMK contributes to Akt-mediated regulation of glucose production in the liver. Furthermore, a loss of IPMK in hepatocytes led to an insulin resistance, which was confirmed by a decrease in insulin-mediated Akt-FoxO1 signaling as well as elevated postprandial glucose level further suggesting that IPMK may be a physiologic regulator of insulin-mediated suppression of glucose production in the liver.

Since the whole body IPMK KO mice are embryonic lethal [1], few studies have investigated the role of IPMK in animal models and the subsequent studies have used tissue-specific IPMK KO models. Recently, Lee et al reported that IPMK deficiency in adipose tissue did not affect energy homeostasis [9]. Hence, to the best of our knowledge, our study is the first to demonstrate a metabolic role of IPMK at the systemic level. Our liver-specific IPMK KO mice showed no difference in body weight or liver weight compared to WT mice under ND-feeding. Interestingly, the increase in body weight and adiposity in male LKO was not due to an increase in food intake but associated with a reduction in energy expenditure. Recently, Guha et al also created liver-specific conditional IPMK KO mice and reported that IPMK mediates transcriptional regulation of autophagy via ULK signaling [28]. It has been shown that the autophagy pathway mediated by ULK1/2 also regulates glucose homeostasis in other cell types [29]. Hence, we cannot exclude the possibility that ULK and other pathways independent of Akt may contribute to the increase in hepatic glucose production in LKO mice.

Our in vivo and in vitro data showed that hepatic IPMK deletion resulted in about 50% of canonical Akt signaling in the liver leading to hepatic insulin resistance. However, we did not find any significant effects of IPMK on body weight or glucose in mice fed a normal diet. These data suggest that IPMK-Akt mediated hepatic insulin resistance may not be sufficient to cause hyperglycemic in the absence of diet-induced obesity. In fact, a previous study showed that the-whole body deletion of Akt2, the predominant form of Akt which accounts for about 85% of total Akt in the liver, did not result in significant glucose intolerance [30]. Recently, there has been increasing appreciation of interaction between genetic and environmental factors, especially diet, in the pathogenesis of diabetes [31; 32]. Chronic exposure to high-calorie diet induces insulin resistance in multiple tissues including liver, muscle, and adipose tissue [33]. Our results showed that HFD-fed LKO mice indeed displayed glucose intolerance and increased gluconeogenesis compared to WT. On an HFD, insulin levels tended to be higher in LKO than WT mice, although the difference was not statistically significant (Supplementary Fig. 4B).

Our data show that insulin-mediated activation of FoxO1 is significantly attenuated in LKO mice. In addition to Foxo1, cAMP response element-binding protein (CREB) is a key transcription factor in hepatic gluconeogenesis [34]. Phosphorylated CREB acts as a scaffold for coactivators such as CBP, p300 and CREB-regulated transcription coactivator 2 (CRTC2) that activate gluconeogenic genes [35; 36]. IPMK directly binds to CREB-binding protein (CBP) and augment its recruitment to promoters of the IEGs in the brain [8; 37]. However, an interaction between IPMK and CBP or physiological functional consequence of this interaction in the liver have not been characterized. A loss of IPMK increased glucose production in the basal as well as insulin-stimulated primary hepatocytes. Further studies are needed to elucidate how IPMK interacts with insulin and other signaling pathways. In conclusion, we have demonstrated that hepatic IPMK plays an important role in hepatic insulin signaling and glucose production. Our study provides a potential therapeutic strategy for treating type 2 diabetes, a condition characterized by insulin resistance and increased hepatic glucose production.

## Abbreviations

IPMK: Inositol polyphosphate multikinase
Akt: protein kinase B
FoxO1: Forkhead box protein O1
Pck1: phosphoenolpyruvate carboxykinase 1
G6pc: glucose 6-phosphatase
PI3K: phosphatidylinositol 3 kinase
HFD: high-fat diet
HGP: hepatic glucose production

## Author contributions

SFK and IJ designed and directed the project. IJ, FA, SJ, and SFK performed the experiment. IJ analyzed data and wrote the first draft of the manuscript. RSA and SFK reviewed and edited the manuscript. All authors approved the paper for submission.

## Acknowledgements

This work was supported by grants from American Heart Association Strategically Funded Research Network (SFRN) Obesity Center (RSA and SFK, 17SFRN33610014) and American Heart Association SFRN fellowship (IJ, 17SFRN33560006).

## Supplemental Materials

**Supplementary Fig 1. IPMK protein levels in tissues.**

(A) Protein levels of IPMK in the indicated tissues from WT mice were analyzed by immunoblotting. (B) Protein levels of IPMK in the liver from ND or HFD-fed WT mice were analyzed by immunoblotting (n = 5 per group).

**Supplementary Fig 2. Effects of hepatic IPMK deficiency in male mice fed normal diet**

(A) Schematic diagram for generation of LKO mice. Protein levels of IPMK in the (B) liver, (C) muscle and (D) adipose tissue from WT and LKO mice were analyzed by immunoblotting (n = 3 per group). (E) Body weights were measured in WT and LKO mice on (WT, n = 5; LKO, n = 8). (F) The ratio of liver weight to total body weight (LW/TW) was measured in WT and LKO mice after overnight fasting (WT; LKO, n = 5) or overnight fasting and 4 hours refeeding (WT; LKO, n = 5). Data are mean ± SEM.

**Supplementary Fig 3. Phenotype of LKO female mice.**

Body weight of WT and LKO female mice fed (A) ND (WT, n = 4; LKO, n = 5) or (B) HFD (WT; LKO, n = 5) was measured weekly from age 8-20 weeks. (C) Intraperitoneal pyruvate tolerance test (PTT) was performed by overnight fasting the mice, and injecting pyruvate (2 g/kg) IP. Tail blood glucose was measured at time 0,15, 30, 60 and 120 minutes (n = 4-5 per group). Data are mean ± SEM.

**Supplementary Fig 4. Glucose and insulin levels in WT and LKO male mice.**

22-week-old male WT and LKO mice fed ND or HFD were euthanized after 6 hours fasting in the morning and blood was collected for measurements of plasma (A) glucose and (B) insulin. Data are mean ± SEM; *p < 0.05, n=10 per group.

