## Supplementary figures for "IPMK modulates insulin-mediated suppression of hepatic glucose production"

### Slide 1
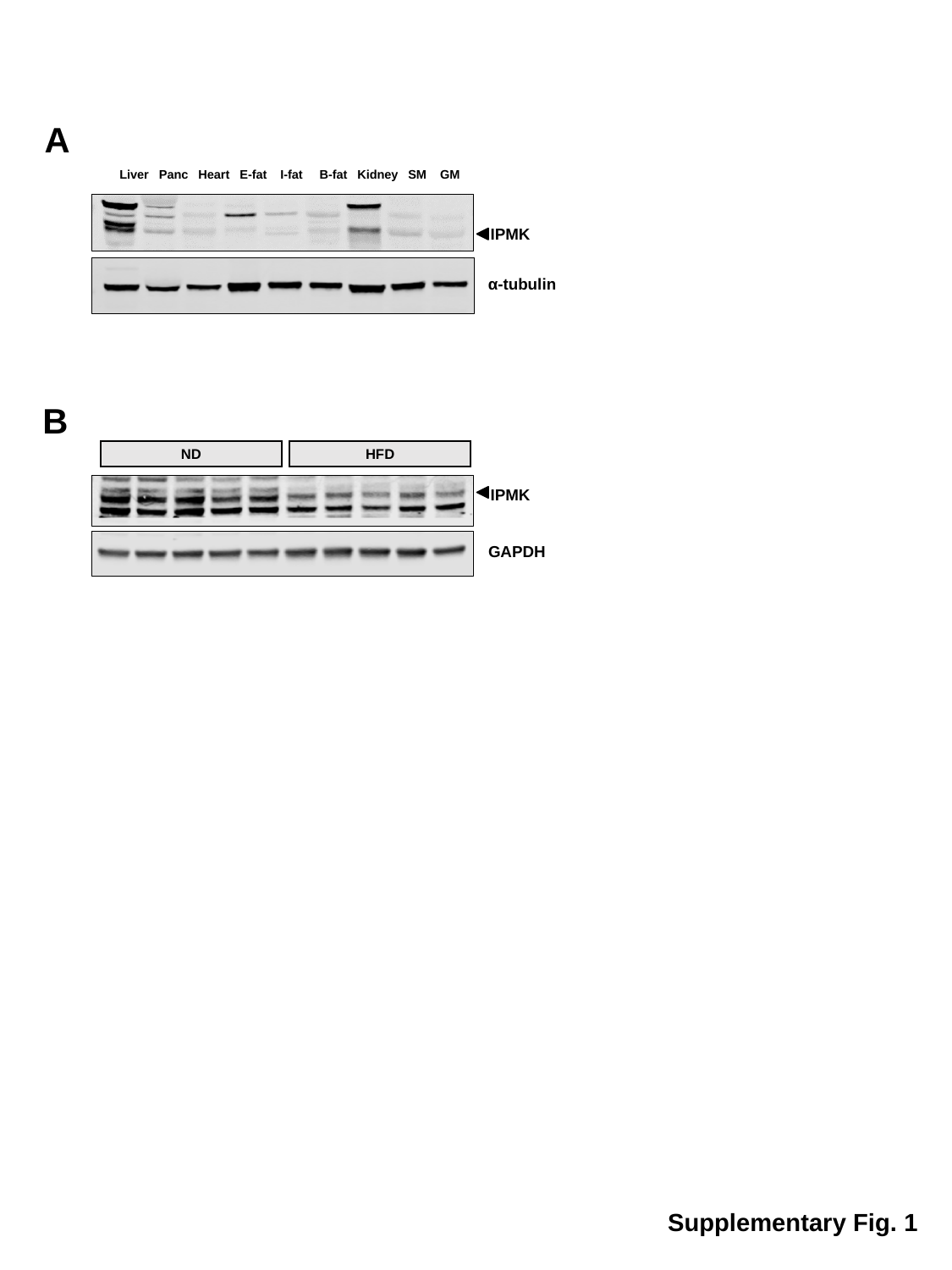

A
Liver Panc Heart E-fat I-fat B-fat Kidney SM GM
IPMK
α-tubulin
B
ND
HFD
IPMK
GAPDH
Supplementary Fig. 1

### Slide 2
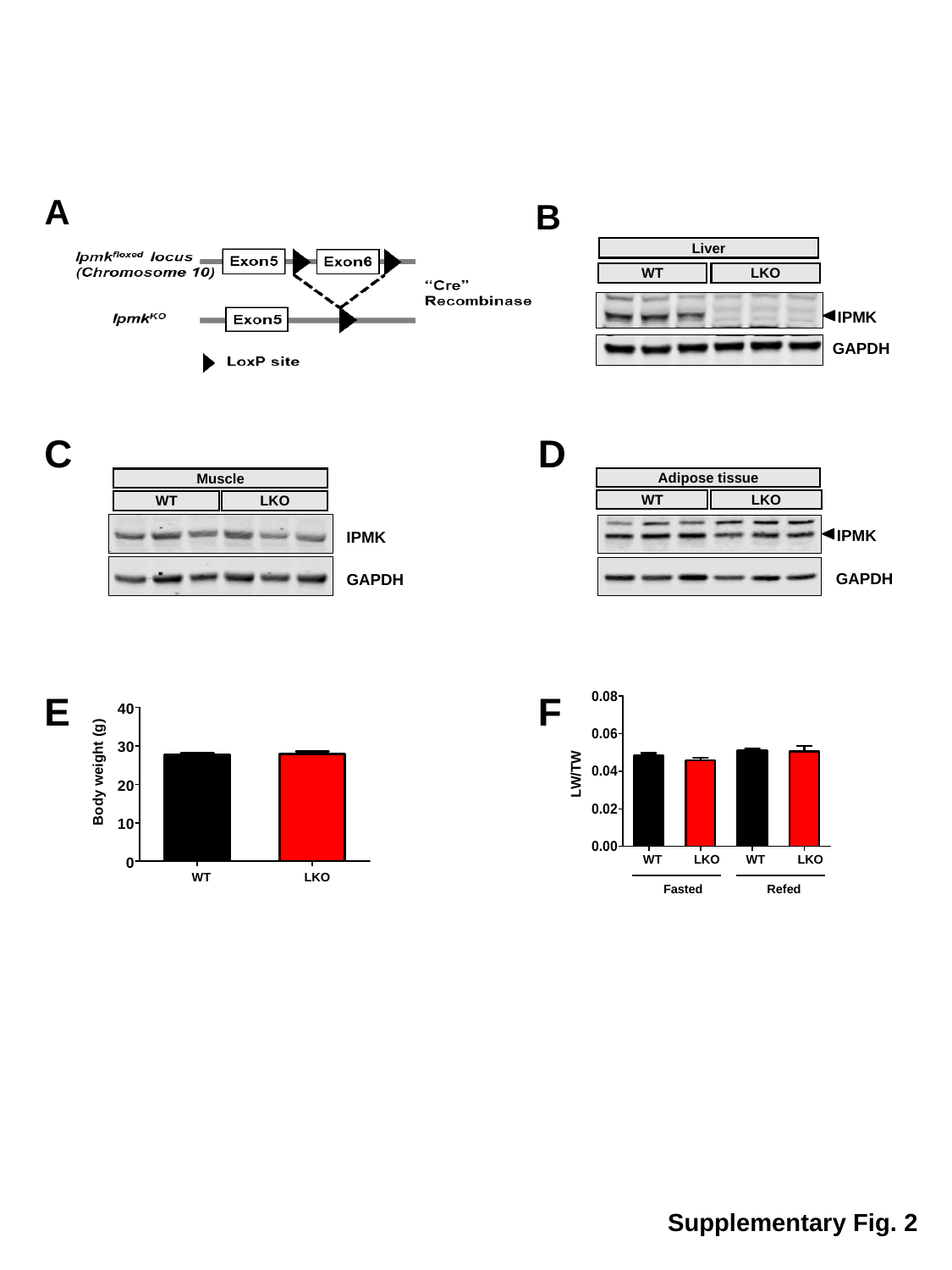

A
B
Liver
WT
LKO
IPMK
GAPDH
C
Muscle
WT
LKO
IPMK
GAPDH
D
Adipose tissue
WT
LKO
IPMK
GAPDH
LW/TW
WT
LKO
WT
LKO
Fasted
Refed
F
E
Body weight (g)
WT
LKO
Supplementary Fig. 2

### Slide 3
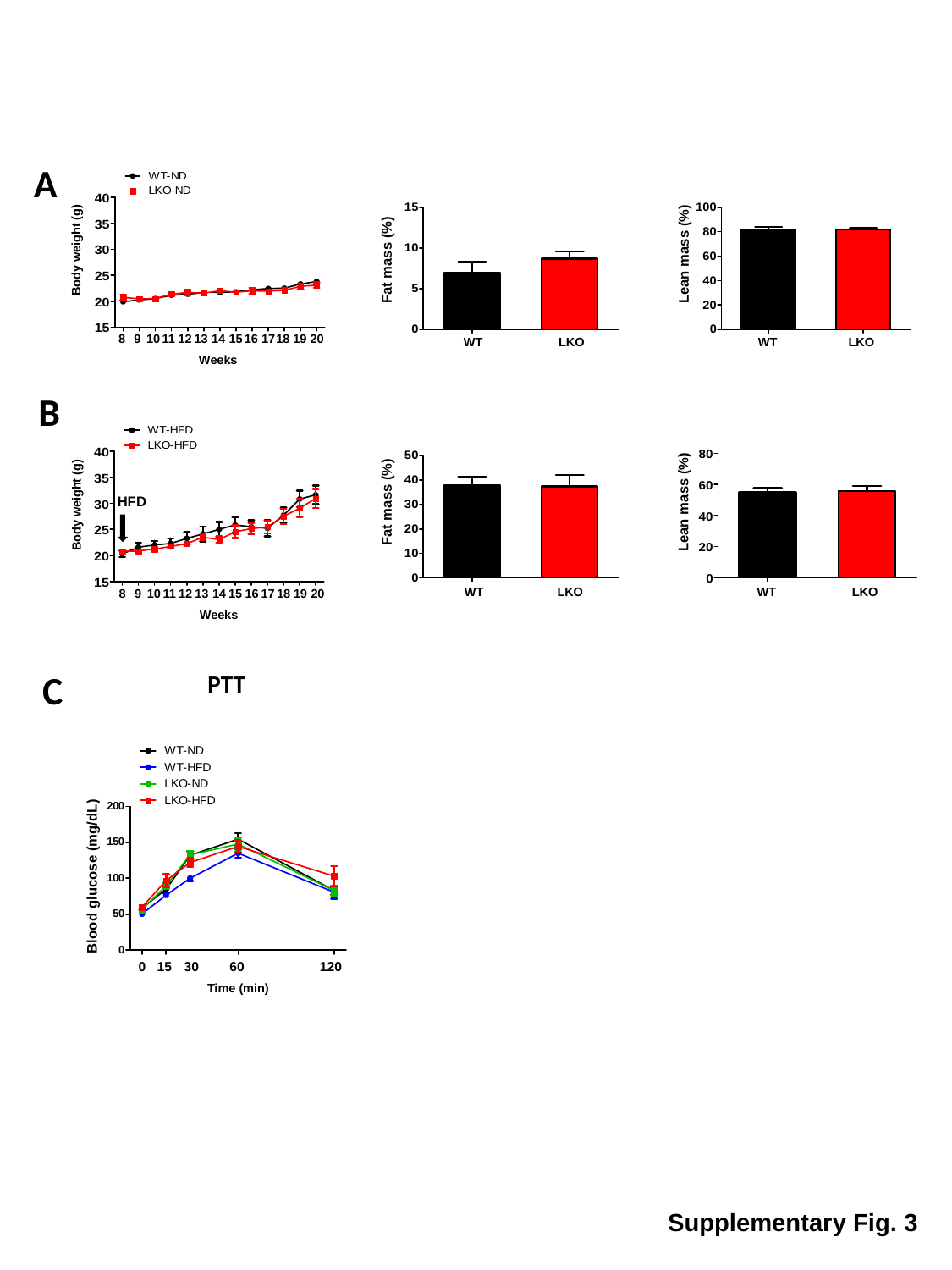

A
Body weight (g)
8
9
10
11
12
13
14
15
16
17
18
19
20
Weeks
Fat mass (%)
WT
LKO
Lean mass (%)
WT
LKO
B
Body weight (g)
8
9
10
11
12
13
14
15
16
17
18
19
20
Weeks
HFD
Fat mass (%)
WT
LKO
Lean mass (%)
WT
LKO
C
PTT
Blood glucose (mg/dL)
0
15
30
60
120
Time (min)
Supplementary Fig. 3

### Slide 4
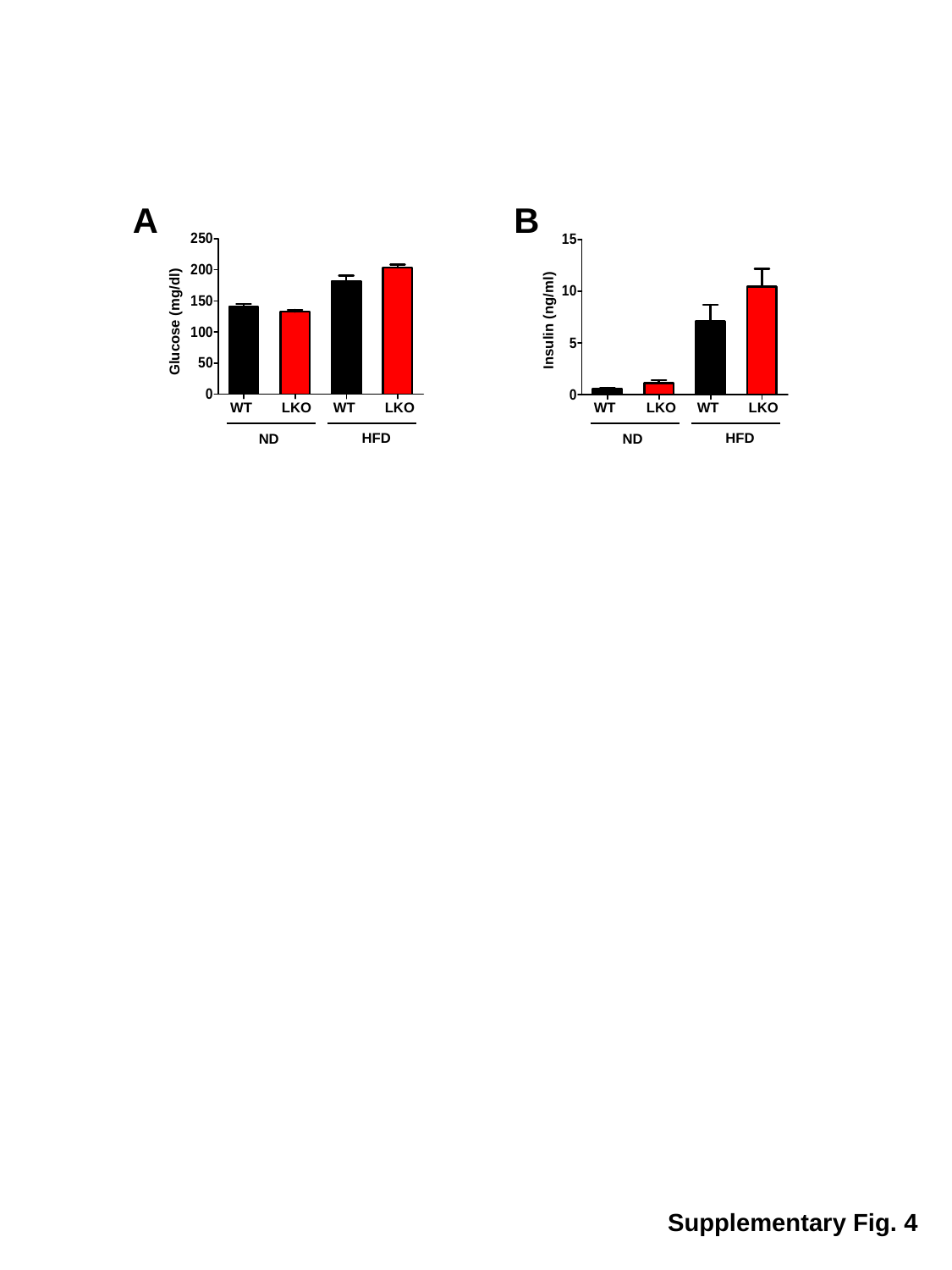

A
Glucose (mg/dl)
WT
LKO
WT
LKO
HFD
ND
B
Insulin (ng/ml)
WT
LKO
WT
LKO
HFD
ND
Supplementary Fig. 4
